# MagNet: Computational Methods for Constructing High-Confidence Protein-Protein Interaction Networks in *Magnaporthe oryzae*

**DOI:** 10.64898/2026.05.11.724438

**Authors:** Hyunbin Kim, Kyeongchae Cheong, Jongbum Jeon, Gobong Choi, Jaeho Koh, Hyeunjeong Song, Yoeguang Hue, Yebin Nam, Byungheon Choi, You-Jin Lim, Jaeyoung Choi, Ki-Tae Kim, Yong-Hwan Lee

## Abstract

*Magnaporthe oryzae*, the rice blast fungus, plays a role as a model organism for molecular plant-microbe interaction research. Studies on the pathogenic mechanism of this fungus revealed many genes involved in signaling pathways. As multi-omics data are being available, genomic-level researches have been conducted to uncover the underlying biological processes during the pathogenesis of *M. oryzae*. Identifying the genome-wide protein-protein interaction (PPI) network is one of the omics-level approaches, which helps to understand signaling and regulatory pathways. However, existing biological network resources of *M. oryzae* are not sufficient to decipher pathogenesis mechanisms due to the abundance of false positives/negatives. In this study, a reliable PPI network database of *M. oryzae*, MagNet, was constructed with three methods, including homology-based ‘Interolog’ search, co-expression network construction, and domain-domain interaction (DDI)-based prediction. With three approaches altogether, the pan-network with 5,600,976 interactions was generated, including 217,531 highly confident interactions supported by all three methods. Experimental data on *M. oryzae* PPIs supported that our PPI network can predict PPIs with higher accuracy compared to the previously constructed databases. MagNet would provide integrated biological network data, which can help to understand the molecular mechanisms of the rice blast fungus. The PPI data can be accessed via https:/magnet.scnu.ac.kr.

## INTRODUCTION

Protein-protein interactions (PPIs) represent fundamental components of cellular processes, as proteins rarely function in isolation. Understanding the complete network of PPIs within cells has emerged as a critical challenge in post-genomic research. While traditional methods of mapping protein interactions, such as yeast two-hybrid assays and bimolecular fluorescence complementation, are valuable, they are insufficient for comprehensively analyzing the millions of potential binary interactions in an organism’s interactome. Consequently, computational approaches have become essential tools for constructing genome-wide PPI networks (Skrabanek et al. 2008).

The computational prediction of interactomes utilizes diverse genomic data sources, including evolutionary relationships, three-dimensional protein structures, genomic positioning, domain architectures, and primary sequence information (Pitre et al. 2008). A prominent computational method is the interaction-ortholog (interolog) approach, which identifies potential interactions based on the evolutionary conservation of protein interactions between homologous pairs across species. This methodology forms the foundation of established databases such as STRING (Szklarczyk et al. 2015). Domain-domain interaction (DDI) analysis represents another significant computational strategy, with databases like DOMINE and IDDI providing valuable insights into protein interaction prediction through domain-level associations (Kim et al. 2012; Yellaboina et al. 2011).

In the context of plant-microbe interactions, PPIs are integral to both pathogen recognition and defense response modulation (Staskawicz et al. 2001). This significance has led to the development of specialized PPI databases for various plant pathogens, including *Fusarium graminearum*, *Magnaporthe oryzae*, and *Ustilaginoidea virens* (He et al. 2008; Zhang et al. 2017; Zhao et al. 2009). Notable applications include the prediction of pathogenicity genes in *F. graminearum* through analysis of PPI sub-networks constructed from verified virulence factors (Lysenko et al. 2013).

*M. oryzae*, the causative agent of rice blast disease, serves as a model organism for investigating plant-microbe interactions due to its significant agricultural impact and experimental tractability (Sadat and Choi 2017). The pathogen’s infection mechanism involves the formation of a specialized structure called the appressorium, whose development and function depend on PPIs within the cAMP-dependent pathway and MAPK cascade signaling (Ebbole 2007; Hamer and Talbot 1998). Following the completion of the *M. oryzae* genome sequence (Dean et al. 2005), research has expanded to encompass multi-omics approaches alongside traditional functional genetics (Franck et al. 2015; Gokce et al. 2012; Kim et al. 2013; Kim et al. 2019). While databases such as MPID and STRING provide predicted PPIs for *M. oryzae*, their reliance solely on interolog-based methods limits the identification of species-specific interactions.

This study presents MagNet, an enhanced PPI network for *M. oryzae* that integrates multiple prediction methods and databases to generate a high-confidence interaction network. The methodology incorporates computational validation and experimental verification from previous studies, enabling systematic analysis through network clustering and integration with existing datasets.

## MATERIALS AND METHODS

### Resources and construction of the PPI network

The study utilized protein sequences and functional annotations of *M. oryzae* strain 70-15, obtained from the Magnaporthe comparative database (Broad Institute of Harvard and MIT, http://www.broadinstitute.org/). Gene Ontology (GO) annotations were acquired from the Gene Ontology Consortium (Meng et al. 2009). The integrated network, designated as MagNet, was constructed using three methodological approaches: homology-based prediction, co-expression analysis, and domain-domain interaction (DDI) prediction (Figure 1). Network visualization and analysis were performed using Cytoscape v3.4.0 (Shannon et al. 2003), with network parameters calculated using the NetworkAnalyzer plugin (Assenov et al. 2008).

**Figure 1.**
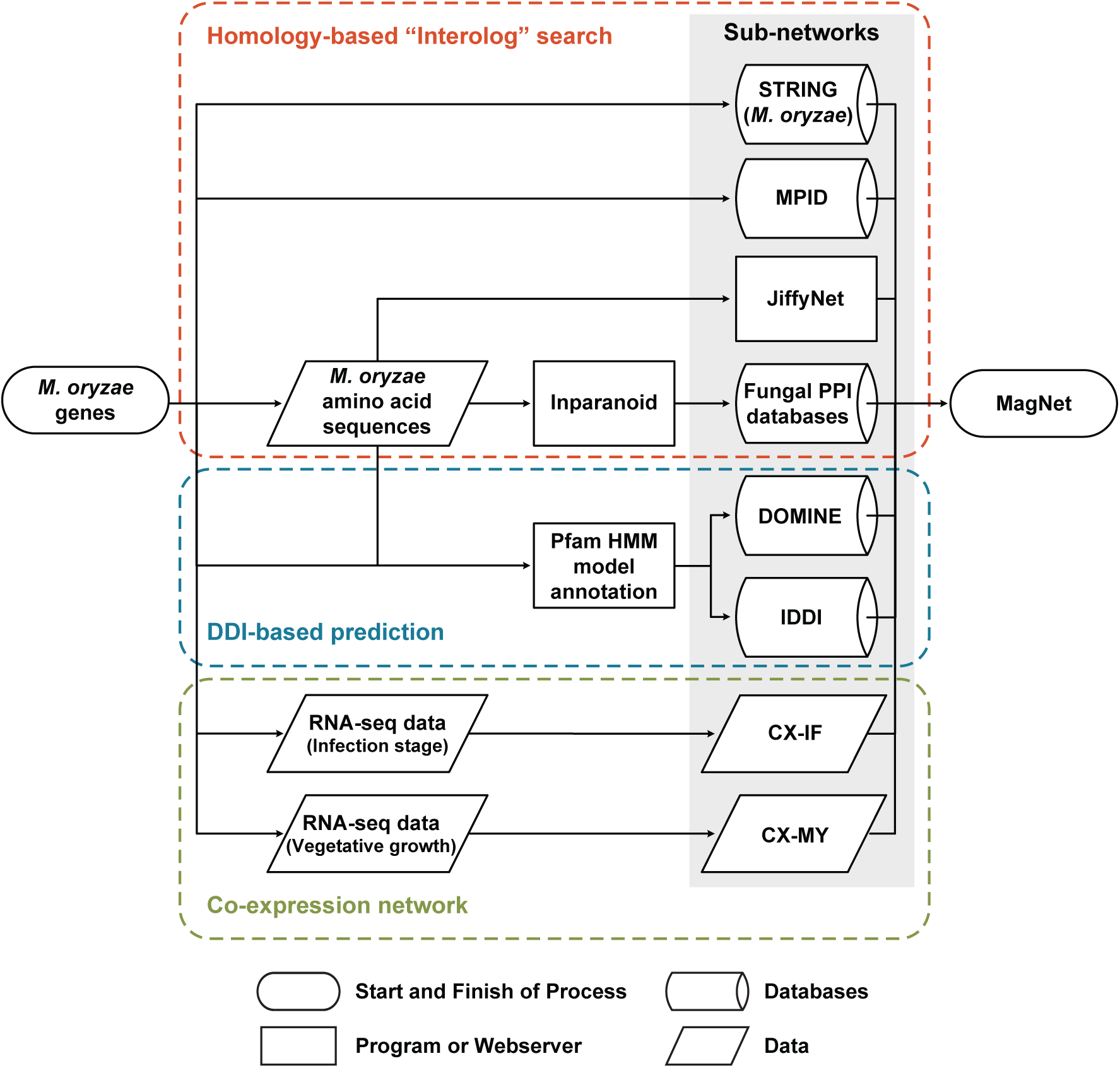
Overview of the construction of multiplex networks of *M. oryzae*. Overview of the three primary methodologies employed in MagNet database construction: homology-based interolog identification, domain-domain interaction (DDI) prediction, and co-expression network analysis. The interolog search utilized Inparanoid 4.0 to identify *M. oryzae* homologs of known interacting protein pairs from fungal PPI databases. DDI-based predictions were generated using domain interaction data from DOMINE and IDDI, applied to *M. oryzae* proteins with Pfam annotations from UniProt. Co-expression networks were constructed using time-series expression data from both infection (CX-IF) and vegetative mycelial growth (CX-MY) stages. The resultant integrated network, MagNet, underwent validation through literature-based evidence and computational simulations.

### Homology-based prediction

The interolog approach was implemented to identify putative interactions between proteins whose homologs demonstrate experimentally validated interactions in other species. Multiple fungal PPI databases were consulted, including DIP, MINT, PINA, INTACT, and BIOGRID (Chatr-Aryamontri et al. 2015; Cowley et al. 2012; Kerrien et al. 2007; Licata et al. 2012; Xenarios et al. 2002). While BioGRID incorporated PPI data from six fungal species (*Aspergillus nidulans*, *Candida albicans*, *Neurospora crassa*, *Saccharomyces cerevisiae*, *Schizosaccharomyces pombe*, and *Ustilago maydis*), other databases primarily contained *S. cerevisiae* interactions. Ortholog identification was performed using Inparanoid v4.1 (Remm et al. 2001), with only perfect-score (1.000) orthologs selected for interolog prediction. Additionally, JiffyNet (Kim et al. 2013), a web-based platform for homology-based PPI network construction, was employed to identify interologs using six template networks: EcoliNet, YeastNet, WormNet, HumanNet, AraNet, and RiceNet (Cho et al. 2014; Hsu et al. 2011; Kim et al. 2015; Kim et al. 2014; Lee et al. 2015b; Lee et al. 2015a). The complete *M. oryzae* proteome (version 8) was analyzed through the JiffyNet web server.

### Domain-domain interaction

Pfam domains were assigned to *M. oryzae* proteins using UniProt data and validated against InterPro v58 (Quevillon et al. 2005). DDI information was extracted from two comprehensive databases: DOMINE and IDDI. DOMINE, previously utilized in *N. crassa* PPI prediction (Wang et al. 2011), encompasses experimentally observed and predicted DDIs from 15 databases. IDDI expands upon this by incorporating ten additional databases, resulting in 26,219 DDIs across 5,140 Pfam domains in DOMINE and 204,705 DDIs across 7,351 Pfam domains in IDDI.

### Co-expression network

Co-expression analysis was employed as a predictive method for identifying potential protein interactions, based on the principle that proteins with similar expression patterns are likely to interact due to shared regulatory mechanisms. Two distinct co-expression networks were generated, corresponding to the infection and vegetative growth stages. The infection stage network incorporated expression profiles from strains KJ201 and 98-06, each comprising five-time points (Dong et al. 2015; Jeon et al. 2020). Comparative co-expression networks were constructed using RNA-Seq data from mycelial growth stages, obtained from the Gene Expression Omnibus (GSM752000, GSM1072034, GSM1375971-3, and GSE51597). Network construction utilized the ExpressionCorrelation plugin in Cytoscape, with Pearson correlation coefficient (PCC) as the co-expression metric. Threshold values were established at 0.903 and 0.846 for infection and vegetative stage networks respectively (P-value < 0.001), with negative correlations excluded from the analysis.

### Validation of predicted PPIs

The reliability of the predicted PPI network was assessed using 97 PPI experimentally validated *M. oryzae* protein interactions collected from 17 published studies (Chen et al. 2008; Choi et al. 2009; Cui et al. 2015; Ding et al. 2010; Jacob et al. 2015; Kulkarni and Dean 2004; Li et al. 2010; Liu et al. 2010; Mehrabi et al. 2008; Park et al. 2006; Qi et al. 2012; Yin et al. 2016; Zhang et al. 2011; Zhang et al. 2014; Zhao et al. 2005; Zhou et al. 2011; Zhou et al. 2012). The validation incorporated data from multiple experimental techniques, including yeast two-hybrid assays, affinity purification, bimolecular fluorescence complementation, and co-immunoprecipitation. These experimental results were compared against predictions from existing databases (MPID and STRING) and MagNet using standard performance metrics (true-positives, true-negatives, false positives, and false negatives).

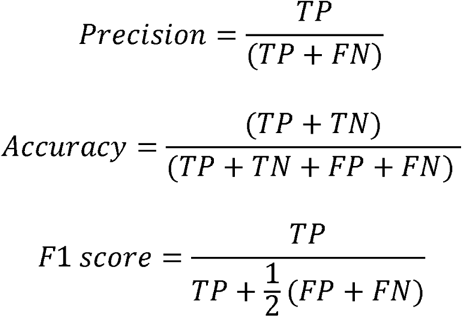

The validation strategy was supplemented with two computational approaches. The first evaluated the reliability of individual prediction methods through comparison with the intersection of MPID and STRING databases. The second analyzed the biological relevance of predicted interactions by examining the shared GO terms between interacting proteins, based on the principle that functionally related proteins are more likely to interact. This analysis compared the GO term depth distribution between MagNet protein pairs and randomly generated pairs.

### Analysis of Interaction Partners

GO term enrichment analysis of putative interaction partners was performed using gProfilerR (Reimand et al. 2007), with significance threshold set at p < 0.05 and false discovery rate correction applied (Meng et al. 2009). Subcellular localization predictions for interaction partners were generated using WolfPSort (Horton et al. 2007).

### Network Topology Analysis

Densely connected network regions were identified using the ClusterONE algorithm (Nepusz et al. 2012), with a significance threshold of p < 0.05. Clusters were annotated using WordCloud summaries of *M. oryzae* gene descriptions and GO annotations, with GO enrichment analysis conducted through the ClueGO Cytoscape plugin (Bindea et al. 2009; Oesper et al. 2011). Hub genes within the high-confidence network were identified using the cytoHubba Cytoscape plugin, which evaluates nodes based on 11 topological parameters (Chin et al. 2014).

### Sub-network analysis

Pathogenic gene information was obtained from PHI-base (Urban et al. 2015), enabling the construction and comparison of pathogenic gene sub-networks across different co-expression conditions. Additional sub-network analysis focused on genes upregulated during appressorium development (Soanes et al. 2012).

A targeted analysis was conducted on MoHox2, a homeobox domain-containing transcription factor essential for conidiation. This analysis examined the sub-network of predicted MoHox2 interactors showing differential expression in Δ*Mohox2* mutants (Kim and Lee 2012). The study also included comparative analysis of interaction partners among related transcription factors MoHox2, MoHox7, and MoHox8, which regulate distinct developmental processes (Kim et al. 2009).

## RESULTS

### Statistics of the integrated network

The integration of multiple prediction methods significantly expanded the coverage of protein-protein interactions compared to existing networks while enabling the extraction of high-confidence interactions. The comprehensive analysis identified 826,128 interolog-based interactions, 3,121,109 domain-domain interaction (DDI)-based predictions, and 1,967,109 co-expression-derived interactions. From the total *M. oryzae* proteome of 12,991 proteins, the coverage analysis revealed that interolog and DDI predictions encompassed 7,138 and 6,226 proteins respectively, while the co-expression network included 10,759 proteins. The final integrated network, comprising all three prediction methods, contained 5,600,976 interactions spanning 11,734 proteins. Network parameters for each constituent database were computed using the NetworkAnalyzer plugin in Cytoscape (Table 1). The complete collection of binary protein interactions has been made publicly accessible through a dedicated database interface at https://magnet.scnu.ac.kr (Figure 2).

**Figure 2.**
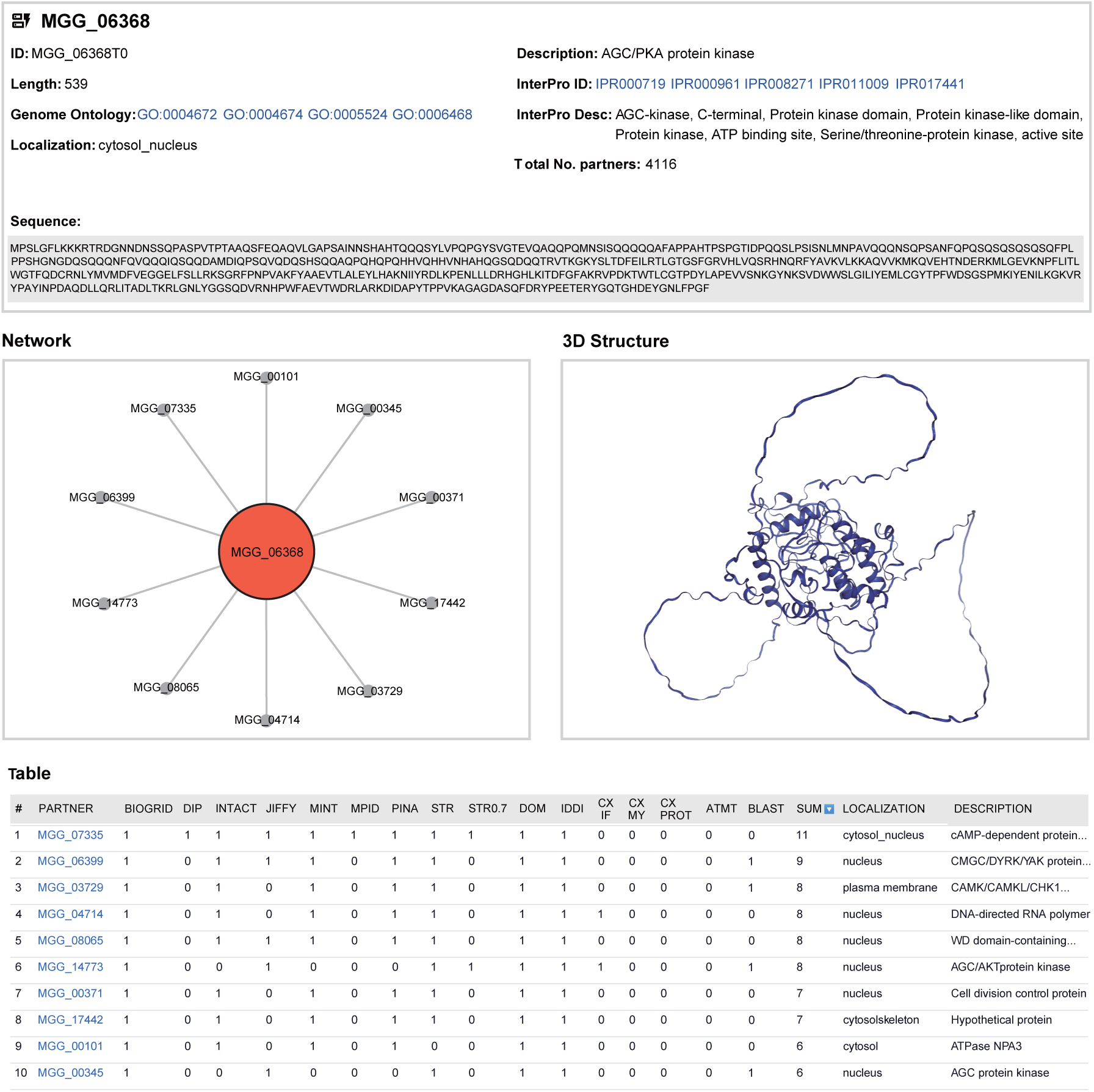
CPKA (MGG_06368) interaction network in MagNet. Network visualization of predicted interaction partners for CPKA (MGG_06368), which encodes the catalytic subunit of cAMP-dependent protein kinase A. The strongest predicted interaction, supported by 11 databases, was with MGG_07335, encoding the regulatory subunit of PKA. Only the 10 most probable entries are presented in the figure.

**Table 1.** Network statistics of MagNet sub-networks.

| Method | Network | Proteins | Interactions | Avg. num<br>of<br>neighbors | Clustering<br>coefficient | Network<br>centralization | Connected<br>components | Network<br>diameter | Network<br>density |
| --- | --- | --- | --- | --- | --- | --- | --- | --- | --- |
| Homology based<br>network | STRING | 6,498 | 1,097,966 | 168.97 | 0.309 | 0.445 | 71 | 9 | 0.026 |
|  | MPID | 3,017 | 11,674 | 7.234 | 0.105 | 0.041 | 108 | 9 | 0.002 |
|  | BioGRID | 3,503 | 166,492 | 61.961 | 0.207 | 0.375 | 4 | 5 | 0.018 |
|  | DIP | 2,344 | 10,110 | 8.438 | 0.130 | 0.054 | 28 | 9 | 0.004 |
|  | MINT | 2,542 | 23,868 | 8.094 | 0.127 | 0.045 | 30 | 10 | 0.003 |
|  | PINA | 2,849 | 38,336 | 26.688 | 0.251 | 0.493 | 2 | 6 | 0.009 |
|  | INTACT | 2,825 | 31,756 | 21.848 | 0.239 | 0.543 | 5 | 6 | 0.008 |
|  | JiffyNet | 4,690 | 266,409 | 113.607 | 0.303 | 0.184 | 2 | 7 | 0.024 |
| Co-expression<br>network | Vegetative stage | 10,356 | 1,173,767 | 226.683 | 0.546 | 0.075 | 6 | 15 | 0.022 |
|  | Infection stage | 8,131 | 819,480 | 201.569 | 0.539 | 0.071 | 5 | 11 | 0.025 |
| Domain interaction<br>network | DOMINE | 5,727 | 1,450,218 | 372.438 | 0.663 | 0.412 | 182 | 10 | 0.065 |
|  | IDDI | 6,231 | 4,247,425 | 991.802 | 0.662 | 0.514 | 51 | 6 | 0.159 |
| High-confidence | MagNet | 6,005 | 217,531 | 71.936 | 0.324 | 0.223 | 57 | 11 | 0.012 |
| Total | MagNet | 11,734 | 5,600,976 |  |  |  |  |  |  |

### High-confidence network extracted from the integrated network

A subset of high-confidence interactions was extracted from the complete predicted interactome to identify reliable interaction partners suitable for experimental validation and hypothesis generation (Figure 3). Validation against experimental evidence demonstrated superior precision of the MagNet high-confidence network compared to existing databases MPID and STRING (Table 2; Supplementary Table 1). The integrated MagNet achieved the highest validation metrics among tested networks, with an accuracy of 0.474 and F1 score of 0.628 (Table 2). Among MagNet’s 43 true positive predictions, eight were unique to MagNet, comprising six interactions identified through IDDI and two through infection stage co-expression analysis. Analysis of MagNet-specific false positives revealed five predictions from IDDI and one from JiffyNet. Experimental validation demonstrated that true positive interactions were supported by an average of 3.88 resources, compared to 2.54 for false positive interactions. Based on these findings, high-confidence interactions were defined as those supported by at least three independent resources. The resulting high-confidence network encompasses 217,531 interactions among 6,005 *M. oryzae* proteins, with an average of 72 interaction partners per protein.

**Figure 3.**
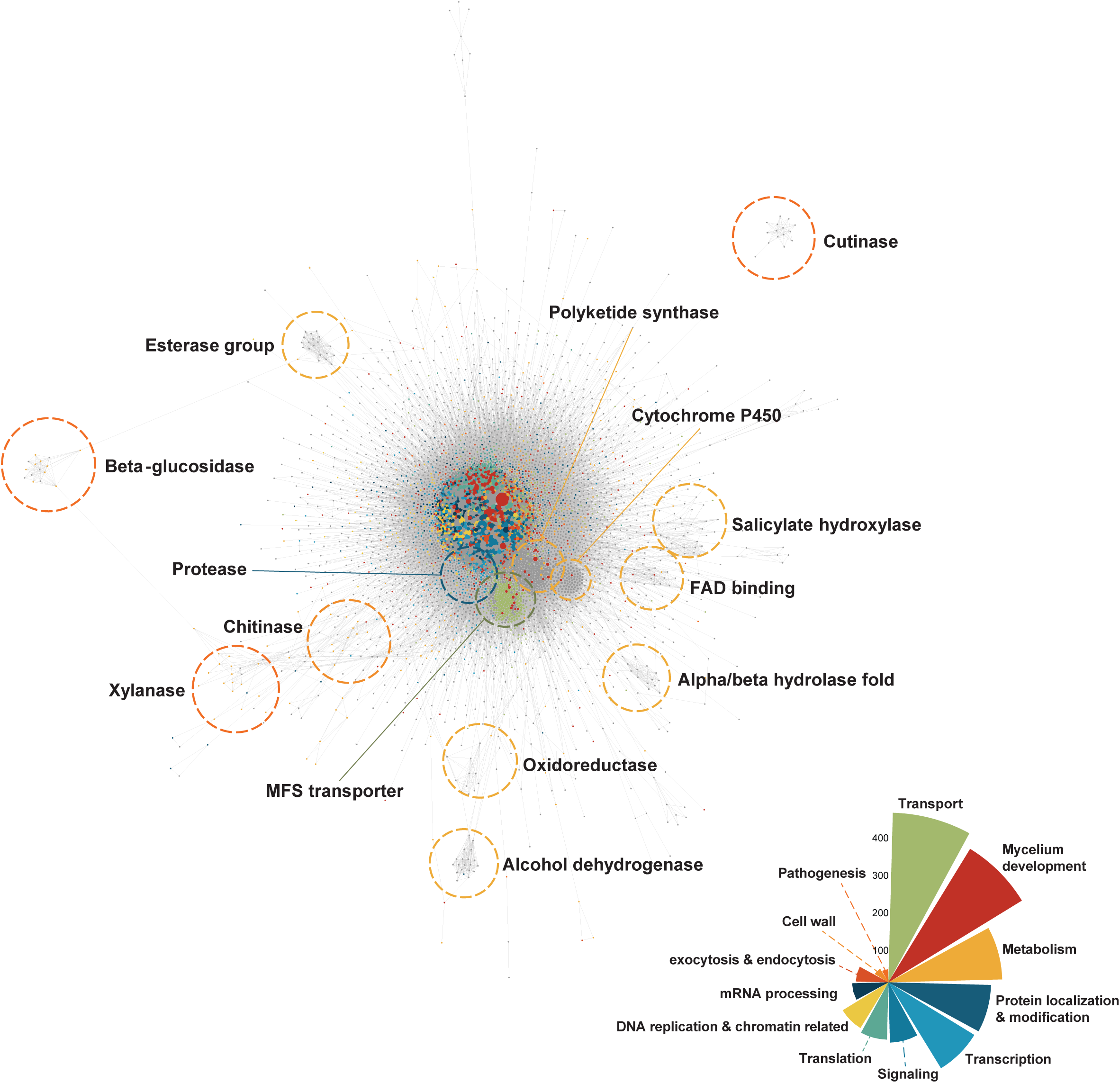
Global architecture of the *M. oryzae* high-confidence network. Comprehensive visualization of high-confidence interactions predicted by three or more independent resources. The network was rendered using Cytoscape 3.4.0, with nodes representing proteins and edges indicating interactions. Node color reflects biological processes based on GO term assignments, while node size corresponds to interaction degree. Edge-dense regions were identified using the ClusterONE algorithm, with clusters annotated based on GO terms and gene descriptions using WordCloud. Notable clusters include MFS transporters, Cytochrome P450, and various enzymatic complexes.

**Table 2.** Validation with literature evidence.

| **Parameter** | **MPID** | **STRING** | **MagNet** | **MagNet_over3** |
| --- | --- | --- | --- | --- |
| True positive (No.) | 7 | 34 | 43 | 26 |
| True negative (No.) | 14 | 9 | 3 | 11 |
| False positive (No.) | 0 | 5 | 11 | 3 |
| False negative (No.) | 76 | 49 | 40 | 57 |
| Precision | 1.000 | 0.872 | 0.796 | 0.897 |
| Accuracy | 0.216 | 0.443 | 0.474 | 0.381 |
| F1 Score | 0.156 | 0.557 | 0.628 | 0.464 |

No false positives were found in MPID, which appears to be due to the small interaction volume of MPID. Integrated MagNet had the most number of true positives and false positives, which led to the highest accuracy and F1 score.

Topological analysis revealed that MagNet exhibits characteristics typical of biological networks, displaying power-law degree distribution with hub nodes showing higher-than-average connectivity (Supplementary Figure 1). This architecture parallels observations in yeast networks, where a small number of highly connected hubs link numerous low-degree nodes (Albert 2005). The network demonstrated a higher average clustering coefficient (0.324) compared to random networks (0.012), aligning with STRING (0.309) but differing from MPID (0.105). Analysis of node degree identified the twenty most highly connected *M. oryzae* proteins (Table 3). These hub proteins predominantly function in post-translational modification, including polyubiquitin, heat shock proteins, and kinases. Notably, hub proteins exhibited elevated expression levels relative to the average *M. oryzae* proteome, suggesting their fundamental importance to cellular function.

**Table 3.** Twenty most highly connected *M. oryzae* protein interaction hubs in the high-confidence network.

| **Locus** | **Protein description** | **Degree** |
| --- | --- | --- |
| MGG_06958 | hsp70-like protein | 1408 |
| MGG_01282 | polyubiquitin | 1137 |
| MGG_11513 | heat shock protein SSB1 | 1108 |
| MGG_01362 | CMGC/CDK/CDC2 protein kinase | 939 |
| MGG_06759 | heat shock protein 90 | 872 |
| MGG_04660 | CMGC/CDK/CDK5 protein kinase | 749 |
| MGG_06044 | ubiquitin-60S ribosomal protein L40 | 741 |
| MGG_07928 | ubiquitin-40S ribosomal protein S27a | 708 |
| MGG_01656 | UV excision repair protein Rad23 | 697 |
| MGG_04719 | guanine nucleotide-binding protein subunit beta-like protein | 679 |
| MGG_01318 | deubiquitination-protection protein dph1 | 669 |
| MGG_09960 | PLK protein kinase | 616 |
| MGG_04905 | CMGC/CDK protein kinase | 608 |
| MGG_06320 | STE/STE20/PAKA protein kinase | 600 |
| MGG_04943 | CMGC/MAPK protein kinase | 599 |
| MGG_14620 | CAMK/CAMKL protein kinase | 591 |
| MGG_14773 | AGC/AKT protein kinase | 582 |
| MGG_00803 | CAMK/CAMKL/AMPK protein kinase | 580 |
| MGG_02503 | glucose-regulated protein | 576 |
| MGG_09887 | NEDD8 | 576 |

### Validation of MagNet with computational methods

The predictive accuracy of individual methodologies incorporated in MagNet was assessed using the intersection of MPID and STRING networks as a validation dataset (Figure 4A). Analysis revealed superior classification performance for BioGRID and IDDI predictions, while co-expression networks, JiffyNet, and DOMINE demonstrated lower concordance with the validation set. Notably, the integration of multiple prediction methods yielded enhanced classification accuracy compared to individual approaches.

**Figure 4.**
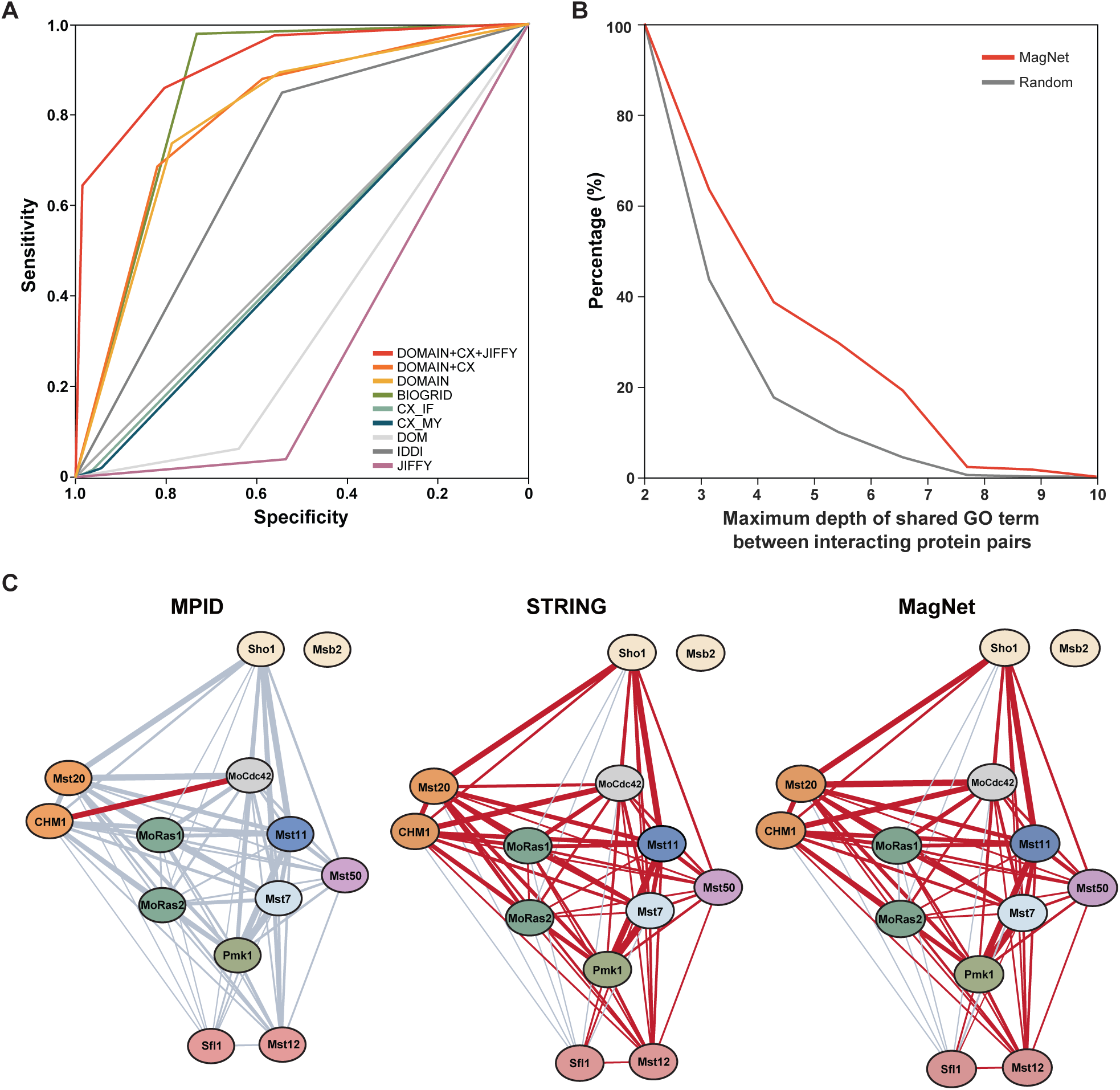
Computational validation of MagNet. **(A)** Performance evaluation of individual MagNet resources using receiver operating characteristic curves, based on comparison with the MPID-STRING intersection set. **(B)** Network validation through analysis of shared GO term depth distributions between predicted and random protein pairs. **(C)** Comparative analysis of MAPK pathway coverage across networks, with red lines indicating interactions predicted by multiple methods. Despite fewer total edges, MagNet maintained comparable pathway coverage to STRING.

Biological validation was performed based on the principle that interacting proteins typically participate in related biological processes and therefore share specific GO terms. Comparative analysis of GO term depth between MagNet-predicted protein pairs and randomly generated pairs demonstrated that MagNet interactions exhibited significantly greater GO term specificity (Figure 4B).

Network coverage analysis was conducted by comparing interaction representation in established biological pathways across MPID (11,674 interactions), STRING (1,097,966 interactions), complete MagNet (5,600,976 interactions), and high-confidence MagNet (217,531 interactions). Despite containing fewer total interactions than STRING, the high-confidence MagNet maintained comparable coverage of established pathways, as evidenced by similar interaction counts in the MAPK pathway (STRING: 56 interactions; MagNet: 55 interactions) (Figure 4C). This pattern of preserved pathway coverage was further corroborated by analysis of the septin complex (Supplementary Figure 2).

### Co-expression network comparison

Comparative analysis of co-expression networks revealed distinct stage-specific protein expression patterns, identifying 2,502 proteins specific to the vegetative stage and 696 proteins characteristic of the infection stage. GO enrichment analysis of vegetative stage-specific proteins highlighted processes including cellular macromolecule metabolism, protein metabolism, cellular component organization, mycelium development, and translation. In contrast, infection stage-specific proteins showed enrichment for terms related to host interaction via extracellular compounds and cAMP response pathways.

Network clustering analysis using the ClusterONE algorithm demonstrated clear differentiation between infection and vegetative stage protein co-expression patterns. The analysis identified 16 statistically significant clusters in the infection stage network and 37 clusters in the vegetative stage network. GO enrichment analysis using gProfileR successfully annotated 10 infection stage clusters and 11 vegetative stage clusters.

Vegetative growth clusters exhibited enrichment for biological processes including rRNA processing, oxidation-reduction reactions, multicellular/mycelium development, transmembrane transport, and DNA replication. Conversely, infection stage clusters were characterized by processes such as translation, protein glycosylation, protein phosphorylation, and single-organism development. This distinct clustering pattern underscores the stage-specific molecular processes governing *M. oryzae* development and pathogenicity.

### Pathogenic gene network

Analysis of genes cataloged in PHI-base revealed distinct network connectivity patterns associated with pathogenicity phenotypes. Statistical analysis demonstrated that genes associated with complete loss of pathogenicity exhibited significantly higher network degrees compared to those with unaffected pathogenicity (t-test, P = 0.03). However, no significant differences in network connectivity were observed between genes associated with complete loss versus reduced pathogenicity, or between reduced versus unaffected pathogenicity phenotypes.

Comparative analysis of co-expression patterns within the pathogenic gene network was performed to evaluate the relationship between infection-stage co-expression and pathogenesis (Figure 5A and 5B). The results demonstrated preferential co-expression of pathogenicity-associated genes during the infection process compared to vegetative growth, supporting the biological relevance of the infection-stage network in understanding pathogenic mechanisms.

**Figure 5.**
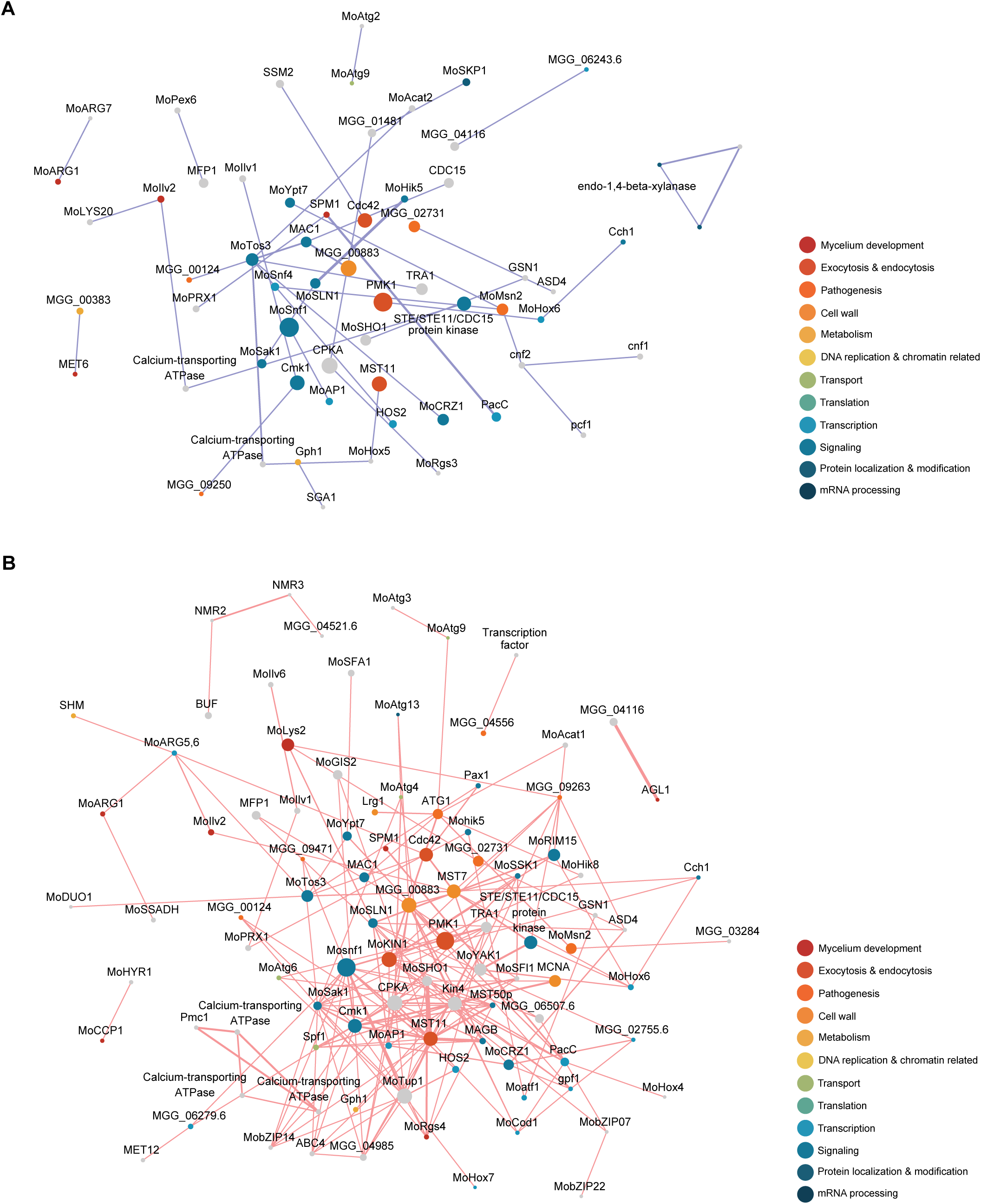
Pathogenic gene co-expression network analysis. **(A)** Co-expression network during vegetative growth was colored blue, and **(B)** the co-expression infection process was colored red. Nodes are colored according to GO biological process annotations. The infection-stage network exhibits enhanced connectivity among pathogenic genes compared to the vegetative growth network.

### Differentially expressed network

Sub-network analysis of differentially expressed genes (DEGs) during appressorium development revealed distinct expression patterns associated with pathogenesis (Figure 6A). Comparison with high-confidence MagNet clusters identified upregulation of functional modules involved in signal transduction, melanin biosynthesis, MFS transport, and cytochrome P450 activity during appressorium formation.

**Figure 6.**
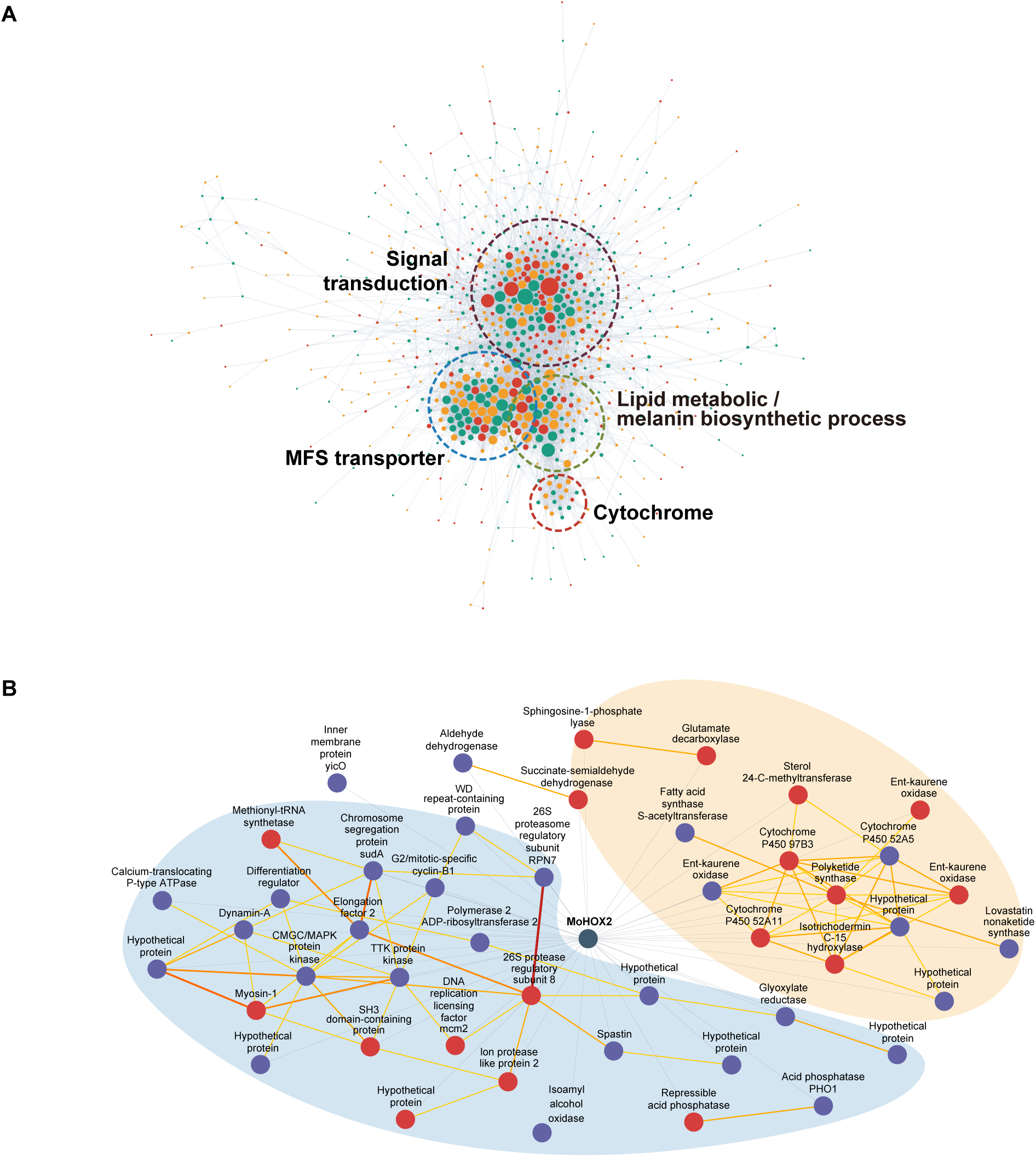
Analysis of differentially up-regulated gene networks. **(A)** Temporal analysis of gene expression during appressorium formation, with nodes colored by expression timing (red: 4-8 hpi, green: 14-16 hpi, yellow: both timepoints). Functional clusters include signal transduction, melanin biosynthesis, cytochrome P450, and MFS transporters. **(B)** Integration of Δ*Mohox2* differential expression data with MoHox2 interaction partners. Blue nodes indicate genes repressed during wild-type conidiation but induced in Δ*Mohox2*, while red nodes show the opposite pattern.

Analysis of the Δ*Mohox2* mutant, which is deficient in conidiation, provided insights into MoHOX2-regulated genes involved in conidial development. Integration of DEG data from Δ*Mohox2* with predicted MoHOX2 interaction partners identified potential regulators of conidiation (Figure 6B). Notable among these were genes encoding kinases, which showed repression by MoHOX2.

Comparative analysis of interaction networks was performed for three HOX transcription factors (MoHOX2, MoHOX7, and MoHOX8), each associated with distinct developmental defects in deletion mutants: Δ*Mohox2* (conidiation deficient), Δ*Mohox7* (appressorium development deficient), and Δ*Mohox8* (impaired invasive growth) (Figure 7). While shared interaction partners among the three HOX proteins were predominantly involved in cellular regulation, MoHOX7-specific partners were enriched for carbohydrate transporters, suggesting a specialized role in appressorium development.

**Figure 7.**
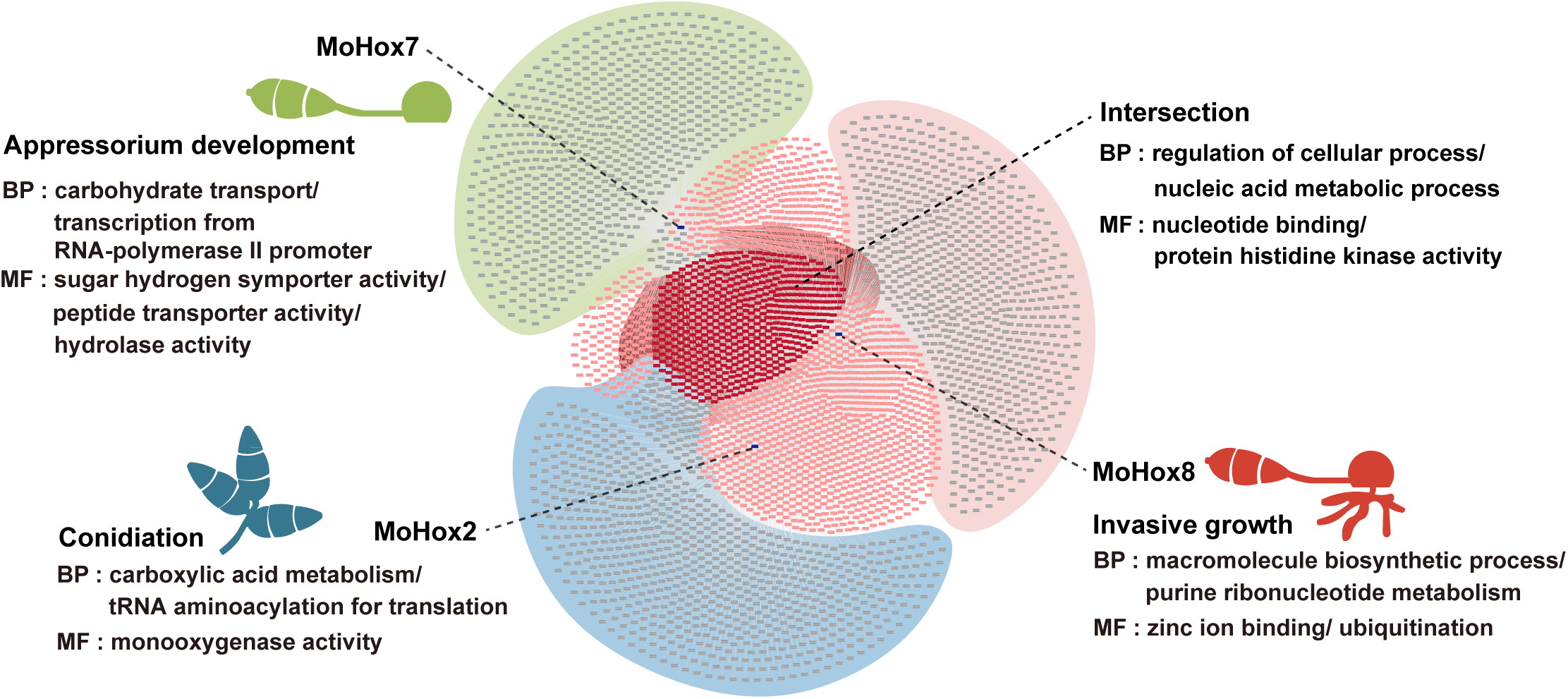
Functional categorization of MoHOX-associated interaction networks in *M. oryzae*. Protein-protein interaction partners of MoHOX2, MoHOX7, and MoHOX8 were analyzed, and GO enrichment was performed to identify significantly overrepresented biological processes (BP) and molecular functions (MF). MoHOX2-associated partners were enriched in processes related to carboxylic acid metabolism and tRNA aminoacylation for translation, indicating a role in conidiogenesis. MoHOX7-specific partners were associated with carbohydrate transport and hydrolase activity, supporting a specialized role in appressorium formation. MoHOX8 partners (red) were enriched in macromolecule biosynthesis and ubiquitination-related functions, consistent with its role in invasive hyphal growth. Intersection nodes represent shared interactors involved in cellular regulation and nucleotide binding.

## DISCUSSION

*M. oryzae* serves as both a critical model organism for studying fungal plant pathogenesis and a significant agricultural pathogen. Given the fundamental role of PPIs in regulating fungal infection processes, comprehensive PPI profiling is essential for understanding disease mechanisms. This study presents MagNet, a high-confidence PPI network containing over 200,000 predicted interactions. Validation through computational methods and experimental evidence demonstrated superior precision compared to existing networks, with MagNet achieving comprehensive pathway coverage despite fewer total edges than STRING.

The integration of diverse prediction methods and updated databases enhanced MagNet’s predictive accuracy. The incorporation of IDDI, absent from MPID and STRING, enabled identification of 43 true positives from 97 experimental validations (Table 2). BioGRID’s extensive repository contributed significantly to true positive predictions in interolog searches. While previous studies of fungal plant pathogens utilized co-expression for validation (Zhang et al. 2017; Zhao et al. 2009), our implementation as a predictive tool identified two novel interactions undetected by other methods, highlighting its value for proteins lacking domain assignments.

This study represents the first integrated application of multiple prediction methods to *M. oryzae* PPI network construction. The clustering analysis revealed functionally associated modules (Ames et al. 2013), enabling identification of protein groups participating in shared biological processes. This approach provides a valuable framework for functional annotation of uncharacterized genes.

Analysis of network hubs aligned with previous observations in yeast studies, where high-degree nodes typically represent essential genes (Krogan et al. 2006). Our findings demonstrated increased network connectivity among pathogenicity-essential genes compared to non-essential genes, supporting this relationship. This network topology enables identification of condition-specific interaction hubs and associated pathways, expanding upon traditional genomic analyses.

Infection-stage co-expression network clustering identified sixteen distinct modules, encompassing processes such as amide biosynthesis, protein modifications, and enzymatic activities. The glycosylation-associated cluster included MoPmt2 (MGG_07190), whose deletion mutant exhibits reduced pathogenicity through impaired cell wall integrity and conidiation (Guo et al. 2016). MoPmt2 clustered with germination regulator MgKIN1 (MGG_01279) and was predicted to interact with cell wall integrity regulator MCK1 (MGG_00883) (Luo et al. 2014; Yin et al. 2016). These associations were further validated through phenotypic correlation analysis using the ATMT database, revealing 19 gene pairs with shared phenotypes in the high-confidence network (Jeon et al. 2007).

The study’s primary limitations include the absence of standardized confidence scoring for prediction methods and its focus on intraspecies rather than host-pathogen interactions. Future development of gold standard datasets and improved secretome characterization would enable more comprehensive prediction capabilities.

MagNet integrates three complementary approaches (interolog prediction, domain-domain interaction analysis, and co-expression network construction) to create a comprehensive PPI network for *M. oryzae*. The identification of pathogenic gene sub-networks and differentially expressed gene networks provides valuable insights into appressorium formation and conidiation pathways. This resource facilitates functional prediction, hypothesis generation, and integration of biological information in *M. oryzae* research.

## Authors’ Contributions

HK, K-TK and Y-HL designed the study. HK, K-TK, JC, and Y-HL wrote the manuscript. KC, YH, YN, BC, K-TK, JC constructed the PPI database. HK, JJ, GC, JK, HS, and Y-JL performed the data mining and analyses.

