## Supplementary Figures for "MagNet: Computational Methods for Constructing High-Confidence Protein-Protein Interaction Networks in *Magnaporthe oryzae*"

### Slide 1
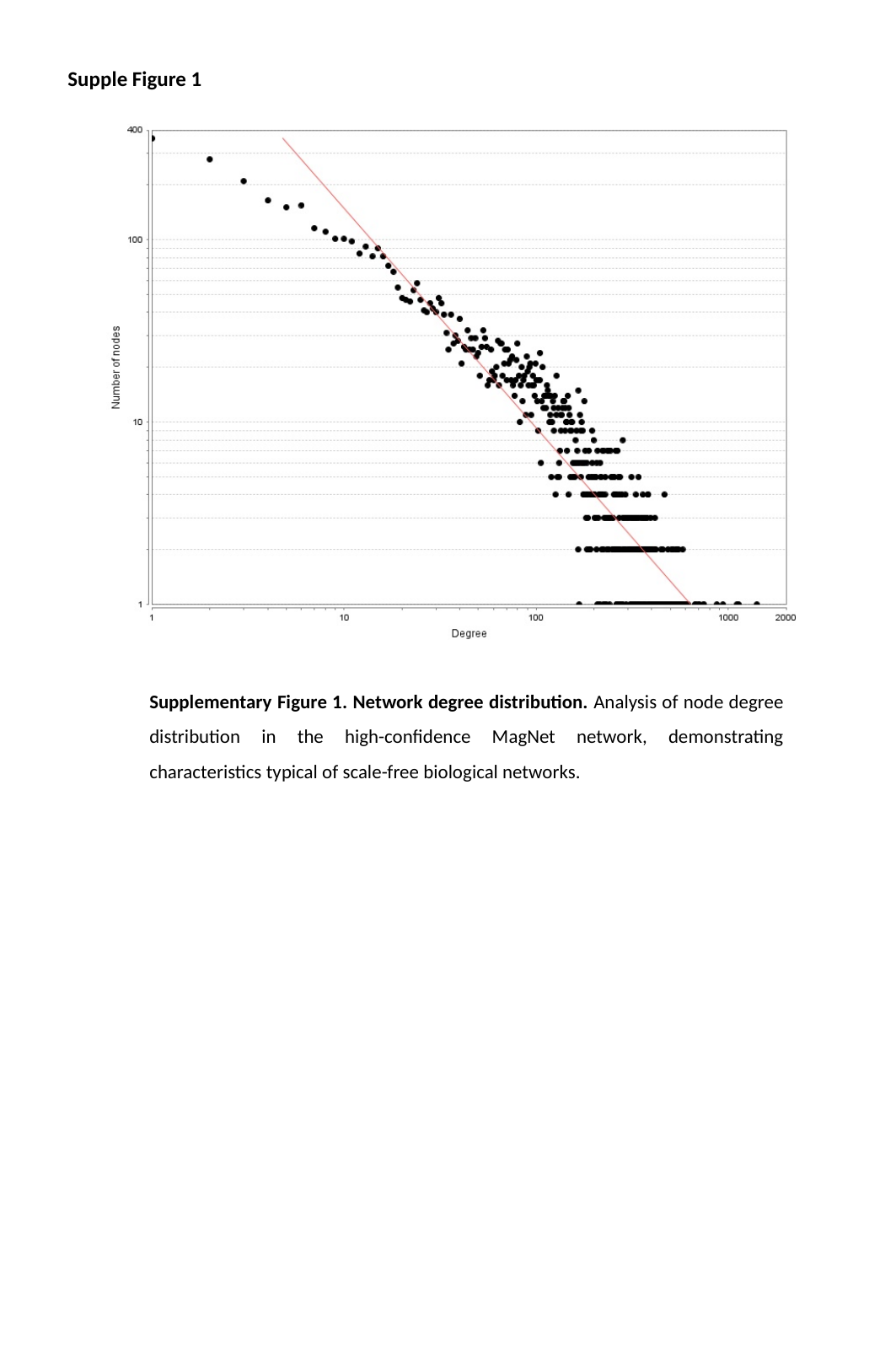

Supple Figure 1
Supplementary Figure 1. Network degree distribution. Analysis of node degree distribution in the high-confidence MagNet network, demonstrating characteristics typical of scale-free biological networks.

### Slide 2
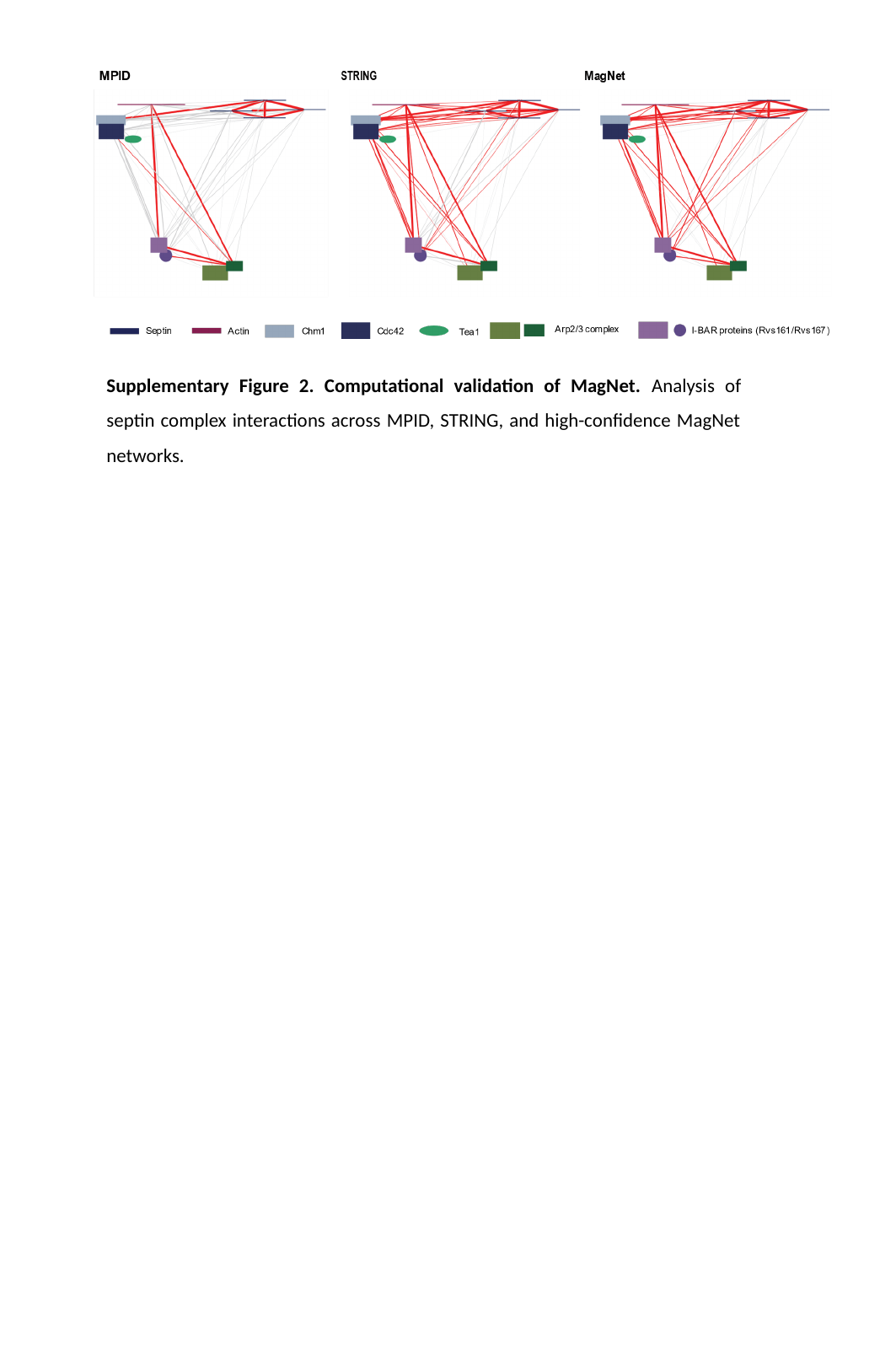

Supplementary Figure 2. Computational validation of MagNet. Analysis of septin complex interactions across MPID, STRING, and high-confidence MagNet networks.
